# Behaviour-based movement cut-off points in 3-year old children comparing wrist- with hip-worn actigraphs MW8 and GT3X

**DOI:** 10.1101/2024.01.06.574473

**Authors:** Daniel Jansson, Rikard Westlander, Jonas Sandlund, Christina E. West, Magnus Domellöf, Katharina Wulff

## Abstract

**Introduction:** Behaviour-based physical intensities have not undergone rigorous calibration in long-term recordings of 3-year-old children’s sleep/activity patterns. This study aimed at (i) calibrating activity counts of motor behaviour measured simultaneously with MotionWatch 8 (MW8) and ActiGraph (GT3X) in 3-year-old children, (ii) documenting movement intensities in 30s-epochs at wrist/hip positions, and (iii) evaluating the accuracy of cut-off agreements between different behavioural activities.

**Methods:** Thirty 3-year-old children of the NorthPop cohort performed six directed behavioural activities individually, each for 8-10 minutes while wearing two pairs of devices at hip and wrist position. Directly observed naturally-occurring behaviours included: watching cartoons, recumbent story listening, sit and handcraft, floor play with toys, engaging in a walk and a sprinting game. Receiver-Operating-Curve classification was applied to determine activity count thresholds and to assign context-guided, physical activity composite classes.

**Results:** Activity counts of MW8 and GT3X pairs of wrist-worn (r = 0.94) and hip-worn (r = 0.79) devices correlated significantly (p < 0.001). Activity counts at hip position were significantly lower compared to those at the wrist position (p < 0.001), irrespective of device type. Sprinting, floorball/walk and floorplay assigned as ‘physically *mobile’* classes achieved outstanding accuracy (AUC >0.9) and two sedentary and a motionless activities assigned into ‘physically *stationary’* classes achieved excellent accuracy (AUC >0.8).

**Conclusion:** This study provides useful cut-offs for physical activity levels of preschool children using two different devices. Contextual information of behaviour is advantageous over intensity classifications only, because interventions reallocate time among behaviours, which allows to establish dose-response relationships between behavioural changes and health outcomes. Our comparative calibration is one step forward to inform behaviour-based public health guidelines for 3-year-old children.

## INTRODUCTION

Movement-related assessments of habitual activities, such as level of physical activity (PA) or timing of sleep/circadian rhythms, have typically taken a segregated rather than integrated approach among research disciplines such as epidemiology, sports medicine, rehabilitation or chronobiology (1–3). Historically, the terminology has also been developed independently, with the term ‘actimeter or actigraph’ in the sleep/chronobiology community and ‘accelerometer’ in the sport/physical activity community (Suppl. Digital Content, **Tab. S1**). The evidence base is similarly segregated, and evidence concerning combinations of movement behaviours that constitute the entire 24-h period using compositional analyses (4) is, to date, rather uncommon, but the interest is growing (5) and studies are emerging (6–8). However, the Commission on Ending Childhood Obesity recognised the importance of interaction among physical activity, sedentary behaviour and adequate sleep on the child’s well-being (9) and instead of focussing on either, physical activity or sleep-wake cycles alone, Canadian and Australian 24-hour movement guidelines were developed (4, 10), which ultimately led to the implementation of WHO guidelines on the amount of *time in a 24-hour day* particularly for children under five years of age (1). Global recommendations, still from a segregated perspective, on the amount of physical activity state that children 3-4 years old should spend at least 180 minutes a day in a variety of physical activities at any intensity, of which at least 60 minutes is moderate- to vigorous-intense physical activity (1). This physical activity recommendation is considered strong, albeit of very low-quality evidence (1).

There are a number of calibration studies for wearable movement sensors using short epochs (5-15s) in preschool children (11–21) (**Tab. S1**). Objective measurements of physical movements with wearables have proven feasible and valid for estimating sedentary time during waking hours in young children in the field (22–24). But comparative studies of valid and reliable devices using longer epochs (30-60s) as in long-term data collection, are limited. In longitudinal recordings over multiple days the distinction of motionless alert from daytime sleep is required and necessitates calibrated actigraphic movement-based sensitivities and sleep algorithms (25, 26).

Equally important, each device type needs to be calibrated for each activity and population (27), epoch as well as commonly used body sites, mainly the wrist and hip (28, 29). The challenges with accelerometer calibration, dividing-up absolute and relative intensities, have been competently outlined by Arvidsson et al 2019 (30) and consensus recently discussed by Migueles et al 2022 (31).

In the segregated tradition, the standard placement of devices for physical activity alone has been at the hip by fitting it on an elastic belt around the waist (32), while in the tradition of sleep and circadian rhythms, devices have typically been attached to the non-dominant wrist (33, 34) **(Tab. S1)**.

Their sensors and firmware, therefore, operate differently according to their purpose: For circadian patterns and sleep timing under free-living conditions, measurements last over complete 24h cycles for several weeks (epoch length: 30 to 60s) with sensors and firmware adjusted for wrist movements (35), while in the physical activity field sensors are usually implemented for high-resolution (seconds), short-term (min-hours) measurements on the hip-position to determine energy expenditure (36–38). For example, activity level outcomes from different devices, such as GeneActiv and ActiGraph GT3X, cannot be considered equivalent because their raw accelerometer data are unequal (39). Compliance and activity prediction is higher for wrist-worn devices and differences according to the machine learning algorithms for classification have been appreciated (32, 39).

In a systematic review on ActiGraph brand devices in children and adults, those worn on the hip, compared to the wrist, were considered more accurate in classifying sedentary behaviour and physical activity in laboratory studies, but not under free-living conditions (40). Recommendation for preschool children were made to attach the device to the *non-dominant* wrist, which also corresponds to the sleep community using actigraphy (41, 42). Although sensors are calibrated to produce equivalent scores between devices of the same model, raw accelerometer output from different models is not equivalent. Scaling factors implemented into algorithms may overcome this inherent discrepancy, provided that the research communities share data collections and meta-data from as many models as possible (2).

There is growing evidence of the influence of ambient light on human physiology and behaviour (43), and therefore, various research-grade data loggers capture not only body movements but also light exposure levels (44). Two of those have been validated for sleep/circadian timing and physical activity in adults and children: MotionWatch 8 (Camntech LTd, UK) and ActiGraph GT3X, both using different algorithms and output modes (45–47). Since we monitor activity in children under free-living conditions in our prospective, population-based NorthPop birth cohort (https://www.katlab.org/ [under people], www.northpop.se), the present study aimed at (i) calibrating movement intensities by determining cut-off points for different behavioural types of activities for MotionWatch 8 and ActiGraph GT3X in 3-year-old children according to an observational protocol, (ii) comparing outcome curves between devices using balanced durations, and (iii) deriving cut-off boundaries for intensities at non-dominant wrist and hip positions. The MW8 epoch of 30 seconds was chosen because this epoch has been validated against polysomnography in simultaneous recordings in adults to derive sleep parameters from movement patterns (48). The GT3X uses the Cole-Kripke (49) and Sadeh et al. (50) algorithms as their standard sleep algorithms. Ultimately, it is intended to assess context-related movements to discriminate time spent ‘*physically mobile*’ (body in motion) from ‘*physically stationary*’ (motionless/sedentary activities, excluding naps/sleep) across a 24h cycle. Motionless alert thresholds will need comparison with existing algorithms defining sleep start/end. We anticipate to be able to assign time spent in behavioural classes in larger cohorts according to cut-off intensity boundaries and, if indicated, to predict individual behavioural classes and compare their allocated time with movement targets for preschool children.

## METHODS

### Participants

In total, 30 children were recruited during the autumn of 2020 from the ongoing prospective, population-based NorthPop cohort. The inclusion criteria were healthy 3-year-old children. Exclusion criteria were any chronic disease or weight outside the normative range (± 2 standard deviations) using a Swedish growth reference (51).

Children in this study were part of a larger actigraphic project within the NorthPop cohort, so all children had experience wearing actigraphs during free-living conditions. Therefore, all participants were already familiar with the procedure, and adding more devices was well-tolerated. Further, all legal guardians received written and oral information about the study and signed a written consent prior to the data collection. The study was conducted in agreement with the declaration of Helsinki and approved by the regional ethical review board (Dnr 2020/01254).

### Procedures

All measurements were conducted between October 2020 and January 2021. Each parent-child pair was studied separately on one test occasion in the E-health laboratory at Umeå University. The E-health laboratory was designed to mimic a small apartment to simulate daily living. We collected the data during daytime between 9.00 to 12.00 or 13.00 to 16.00. Basic anthropometric measurements were collected for each child. Height was measured to the nearest 0.1 cm using a folding rule and body mass to the nearest 0.1 kg using an electronic scale. A pilot evaluation of the feasibility of various activities was carried out before data collection. A final study protocol was developed to monitor three stationary behaviors and three physically demanding behaviors typical for children to engage in at this age under free-living conditions. The six behaviours represent different movement patterns and intensity levels: Physically *mobile* levels endorsed physically ‘vigorous’, ‘moderate’ and ‘light’ activities, and physically *stationary* levels endorsed ‘sedentary crafts’, ‘recumbent listening’ and ‘sedentary screen time’ (**Fig. 1**).

**FIGURE 1.**
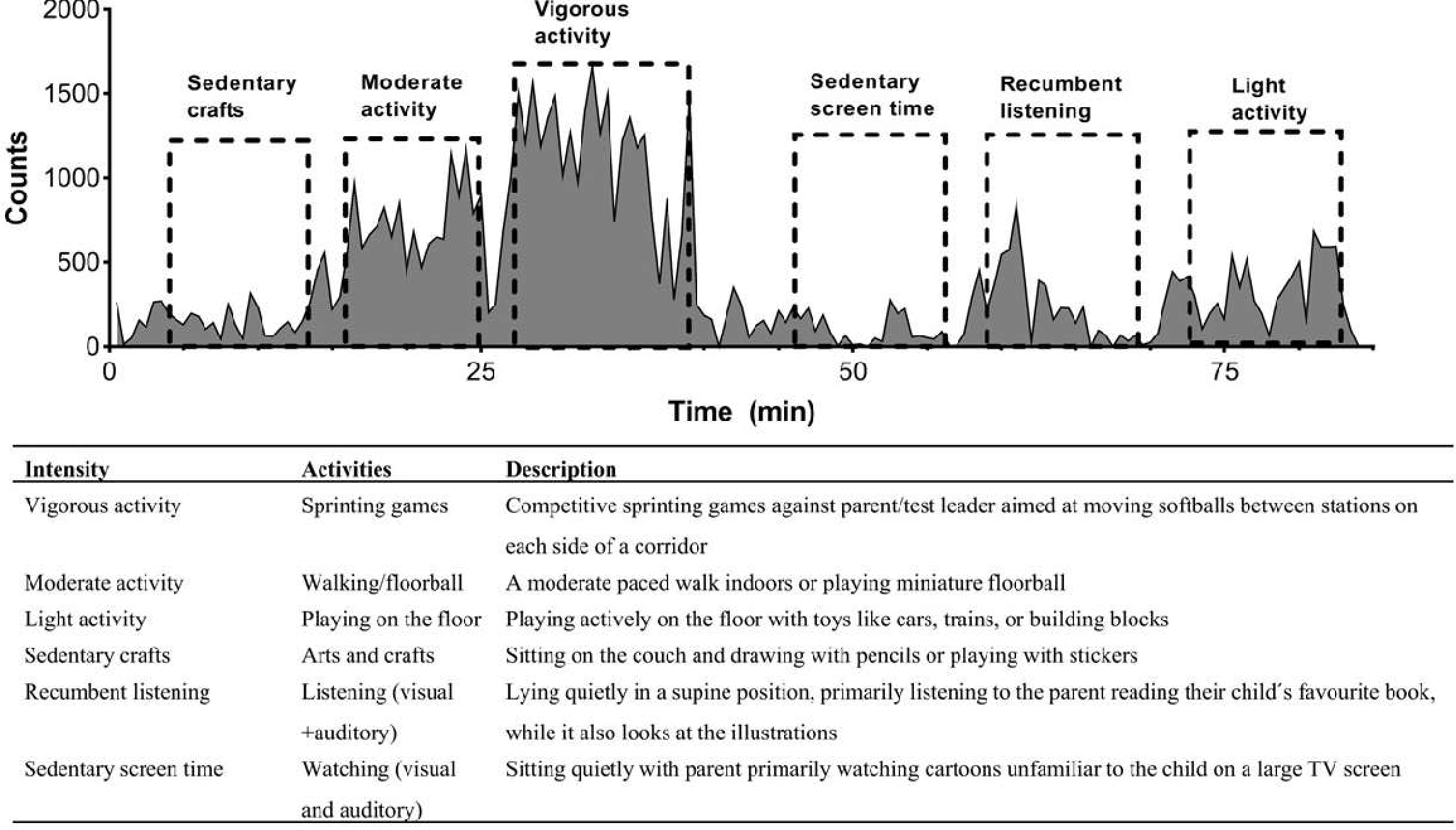
Top: Representative example of an area graph derived from wrist activity patterns of a child performing six investigator-observed behaviours (dashed boxes) shown for a MotionWatch 8 device attached to the non-dominant wrist. The activity intensity was transformed into count values recorded every 30 seconds (epochs). Each type of behaviour was observed by stop-watch to last between 8 to 10 minutes. Bottom: Explanatory table for the six behaviours in terms of terminology referring to intensity, kind of activity and its description.

Sedentary screen time involved watching a cartoon on a large screen TV with their parent and recumbent listening involved a parent reading their child’s favourite story. The ‘sedentary crafts’ activity involved the child sitting at a table, drawing or placing stickers in a sticker book. The three active behaviours were planned to be progressively more intense with light activity, repeesented by playing on the floor with toys, moderate activity by walking or playing floorball, and vigorous activity represented by competitive sprinting games. Each activity was performed for approximately 10 minutes, with a minimum of 3 minutes rest between activities. If the activity was less than 8 minutes, we repeated that specific activity until it was within the time limits (8-10 min). The activities could be executed in any order depending on the child’s preferences, whilst a relaxation break with a snack was offered after about half way through the activities. After the vigorous activity, the child had a little longer break (≈10 min) to minimize the influence of fatigue on the following activity. Two research team members were skilled to explain the study to the child and their parent, to handle the technical devices and software, and trained in momentary time sampling observation in-situ. Each activity was carefully timed and start and stop time was documented.

Children were fitted with four movement sensors (MotionWatch 8 ([MW8], CamNtech Ltd., Cambridge UK), ActiGraph [GT3X], ActiGraph, LLC, Fort Walton Beach, FL) and a heart-rate monitor (Actiheart 5, CamNtech), whose data were not part of the current analyses. A pair of MW8 and GT3X were attached to the non-dominant wrist and another pair to the hip. The hip-worn sensors were mounted to an elastic belt so that the black circle of the GT3X and the label of the MW8 were both facing downwards. The wrist-worn sensors were mounted to the same strap pointing towards the hand, following our standard operational procedures. The children wore the belt around the waist, placed 1-2 cm beneath the umbilicus. The same four devices were used in all children during all activities to minimise variability between sensors. This set-up enabled comparisons between different types of sensors in the same position (wrist-wrist versus hip-hip) as well as within and between sensors across different positions (wrist versus hip).

Data from MW8 and GT3X were captured in 30-second epochs. The GT3X is a lightweight (27 g), relatively obtrusive (3.8 x 3.7 x 1.8 cm) device with a rechargeable battery. Batteries were charged before each trial to avoid missing data. The GT3X collects motion data on three axes (x,y,z) and is designed to record accelerations ranging from 0.05 to 2.5 g. The sampling frequency of 30 Hz and “triaxial mode” was used for both GT3X devices and the “vector magnitude (VM)” for the analyses. The MW8 is battery-powered and weighs 9.1g. It is of relatively unobtrusive dimensions (3.6 x 2.82 x 0.94 cm), which is an advantage for long-term wear in younger children (e.g., 3-year-olds). Its continuous operation has the option of single-axis or tri-axial recording mode (producing a vector magnitude count per epoch). The MW8 has a built-in light sensor and a time-stamp marker button. The MW8 sensing ranges between 0.01 and 8g, and the minimum ‘not moving’ threshold is 0.1g. Data are sampled at 50 Hz and bandwidth-filtered between 3 Hz and 11 Hz. An ‘activity count’ (unitless) is derived from the highest of the 50 samples/second, and these values are accumulated over the length of an epoch (30 seconds). Both MW8 devices were set up to record in single-axis “MotionWatch Mode 1”, and not tri-axial mode, because this mode has been validated against polysomnography. It also allows longer recording periods, as implemented in the ongoing collection of sleep-wake and light exposure data in the Northpop cohort. After data collection, the raw time series data were downloaded using the devices’ respective proprietary software and exported into an Excel spreadsheet. The start and stop times of the behavioural observation protocol were matched to the 8 to 10 minutes-long periods of raw time series data in Excel. Periods of the same behaviour across the children were successively concatenated, separately for each position and device (**Tab. 2**). The total duration per behaviour was calculated and compared between the six behaviours to ensure similar data length distribution (**Fig. S1**).

### Statistical analysis

All statistical analyses were performed using the SPSS statistical package (SPSS, v. 27, Chicago, IL). Anthropometrical data at baseline were compared between males and females using a standard Students Unpaired T-test. The two-tailed Pearson product-moment correlation was used to determine the relationship between the activity measured with the GT3X and MW8. To approximate the similarity in the shape of the data distribution between algorithms, the data of wrist-worn devices were analyzed with linear regression and the data of hip-worn devices with nonlinear regression models. Activity counts were plotted as boxplots in a log scale to visualise their proportional overlap between adjacent behavioural activities.

Receiver Operator Characteristics (ROC) curves were used to determine cut-off points (intensity thresholds) from activity counts of pre-defined, observer-based behaviours (52, 53). ROC curves have previously been used to determine activity count thresholds in children between 4-8 years for Actigraph and MW8 models separately (12, 13, 47). Here we used both brands simultaneously and applied ROC curves to examine the classification accuracy across brands and positions. We used a binarised approach in ROC curve classification acknowledging the activity counts to be of descending order from vigorous PA > moderate PA > light PA > sedentary > motionless alert. Classification can be performed in two ways: pair-wise ‘One-vs-One’ (**Fig. 2a**) or one-group/composite against all other ‘One-vs-Rest’ (**Fig. 2b**). Although we report One-vs-One details in Suppl. Digital Content (**Tab. S2**), here we focus on the One-vs-Rest (OvR) scheme (54). We collapsed counts of certain adjacent behavioural activities (and assumed as one) and compared those against all other collapsed behavioural activities (**Fig. 2b)**. The behavioral activities were assigned with intensity class labels: the most vigorous activity ‘Sprinting’ was assigned to ‘vigorous PA’ (VPA); ‘Sprinting’ and ‘Floor ball’ combined were assigned to ‘moderate-vigorous PA’ (MVPA); ‘Sprinting’, ‘Floor ball’ and ‘Play on floor’ combined were assigned to ‘light-moderate-vigorous PA’ (LMVPA); and all these together were labelled ‘*mobile* PA’ class. They were set against all physically ‘*stationary’* behaviours, thereby dividing ‘*mobile PA*’ from ‘*stationary PA*’. *Stationary* PA included ‘sedentary crafts’, ‘recumbent listening’ and ‘sedentary screentime’. We merged ‘sedentary crafts’ and ‘recumbent listening’ behaviours into ‘sedentary PA’ (SED) on the basis of their considerable overlap in movement patterns. The most immobile behaviour ‘sedentary screentime’ was assigned to ‘motionless alert’ (MOA). We delibertly split off MOA from SED to better understand their closeness to quiet awake and sleep. We did not run a separate ROC-AUC for the SED, but report the range between lower boundary of LMVPA intensity and the upper boundary of MOA intensity.

**FIGURE 2.**
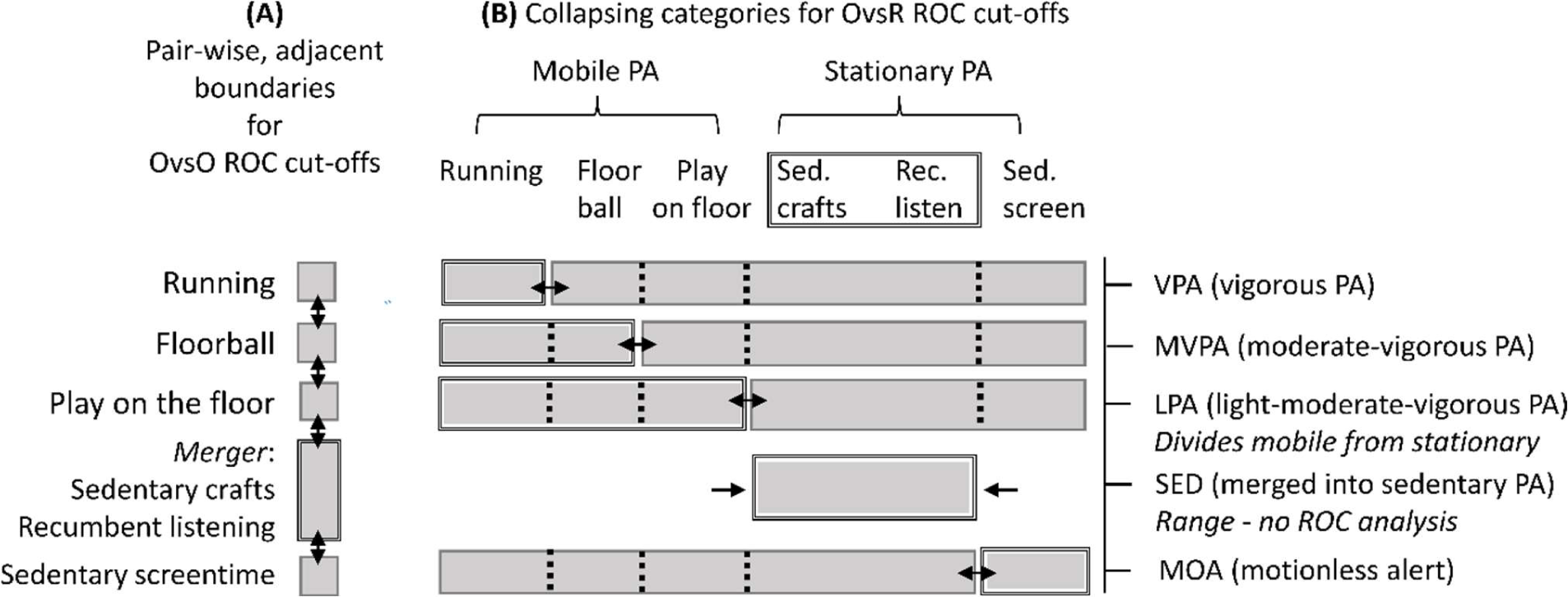
Schematic description of ROC analysis procedures: (a) One versus one (OvsO) and b) one versus the rest (OvsR) in order of descending intensities. Double-sided arrows indicate the cut between categories for each ROC analysis. In OvsR, two sedentary categories (Sedentary craft + Recumbent listening) were merged into SED (for which ROC analysis were done only in OvsO, see **Tab. S2**). Moreover, SED and MOA were collapsed into an composite ‘*stationary* PA’ class and tested against the composite ‘*mobile* PA’.

To determine the best possible compromise between sensitivity and specificity for each cut-off, we applied the Youden Index (*J*), which is determined by calculating the sum of sensitivity and specificity minus one (55). The cut-offs (synonym for intensity thresholds) were derived from the raw data distribution’s highest total sensitivity and specificity ratio. All values are reported as mean ± standard deviation (SD). The Area Under the Curve (AUC) quantified how well the ROC curve performed at classifying data. Values for AUC ranges from 0 to 1. It is used as a measure of accuracy for the ROC curve in discriminating classes, here in classifying count cut-offs from pre-defined, observed behavioural activities. ROC-AUC values above 0.90 are considered outstanding, 0.80-0.89 excellent, 0.70-0.79 acceptable (fair), less than 0.70 are considered poor, and an AUC as low as 0.5 indicates no greater predictive ability than by random guessing (56). A significance level of p < 0.05 was considered significant.

## RESULTS

Anthropometrical data for all participants are presented in **Tab. 1**. The study included 30 participants, 20 males and 10 females. There was no significant difference in age (p = 0.69), height (p = 0.898) or waist circumferences (p = 0.052) between males and females. The males were heavier than females (p = 0.023), which was also reflected in their BMI values (p < 0.05). Mean activity counts were lowest when children were watching cartoons and highest when running in a sprinting game (**Fig. 1** and **Tab. 2**).

**TABLE 1.**
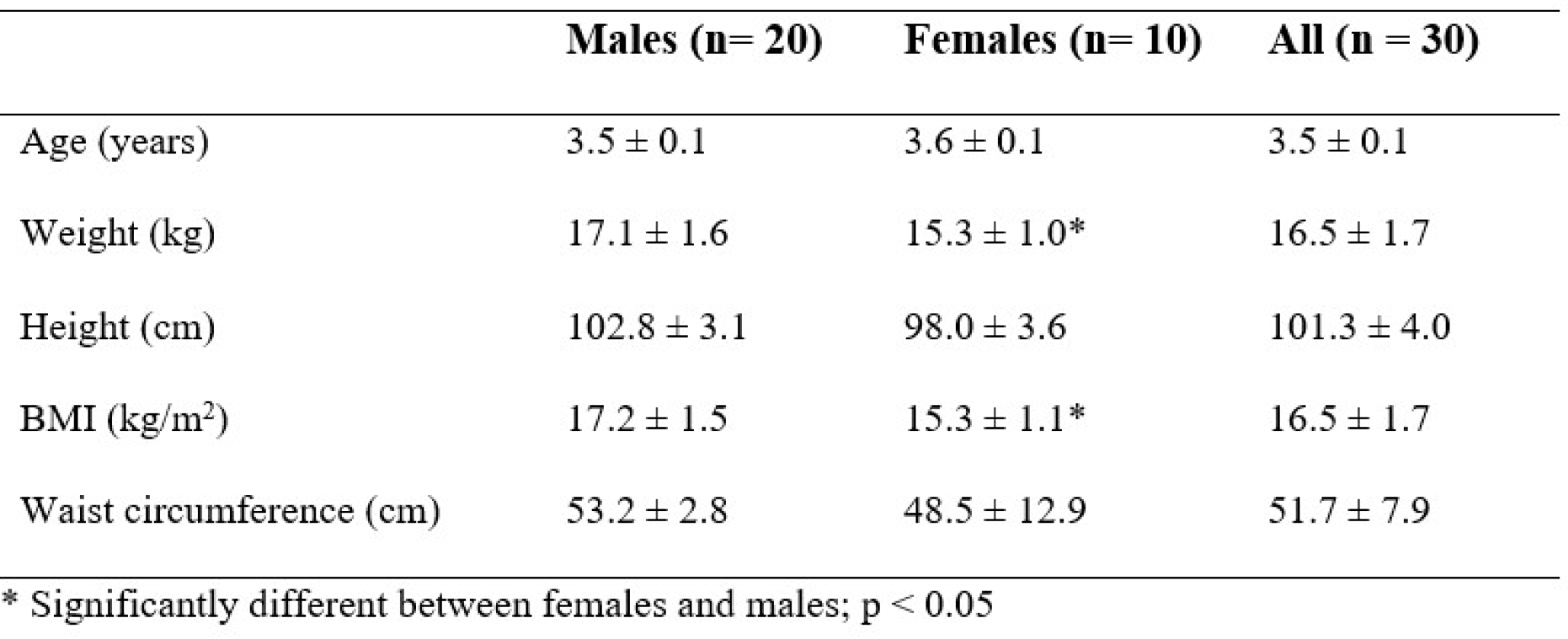
Descriptive characteristics of the participants (n = 30). Shown are means ± SD.

|  | **Males (n= 20)** | **Females (n= 10)** | **All (n = 30)** |
| --- | --- | --- | --- |
| Age (years) | 3.5 ± 0.1 | 3.6 ± 0.1 | 3.5 ± 0.1 |
| Weight (kg) | 17.1 ± 1.6 | 15.3 ± 1.0* | 16.5 ± 1.7 |
| Height (cm) | 102.8 ± 3.1 | 98.0 ± 3.6 | 101.3 ± 4.0 |
| BMI (kg/m<sup>2</sup>) | 17.2 ± 1.5 | 15.3 ± 1.1* | 16.5 ± 1.7 |
| Waist circumference (cm) | 53.2 ± 2.8 | 48.5 ± 12.9 | 51.7 ± 7.9 |

\* Significantly different between females and males; p < 0.05

**TABLE 2.**
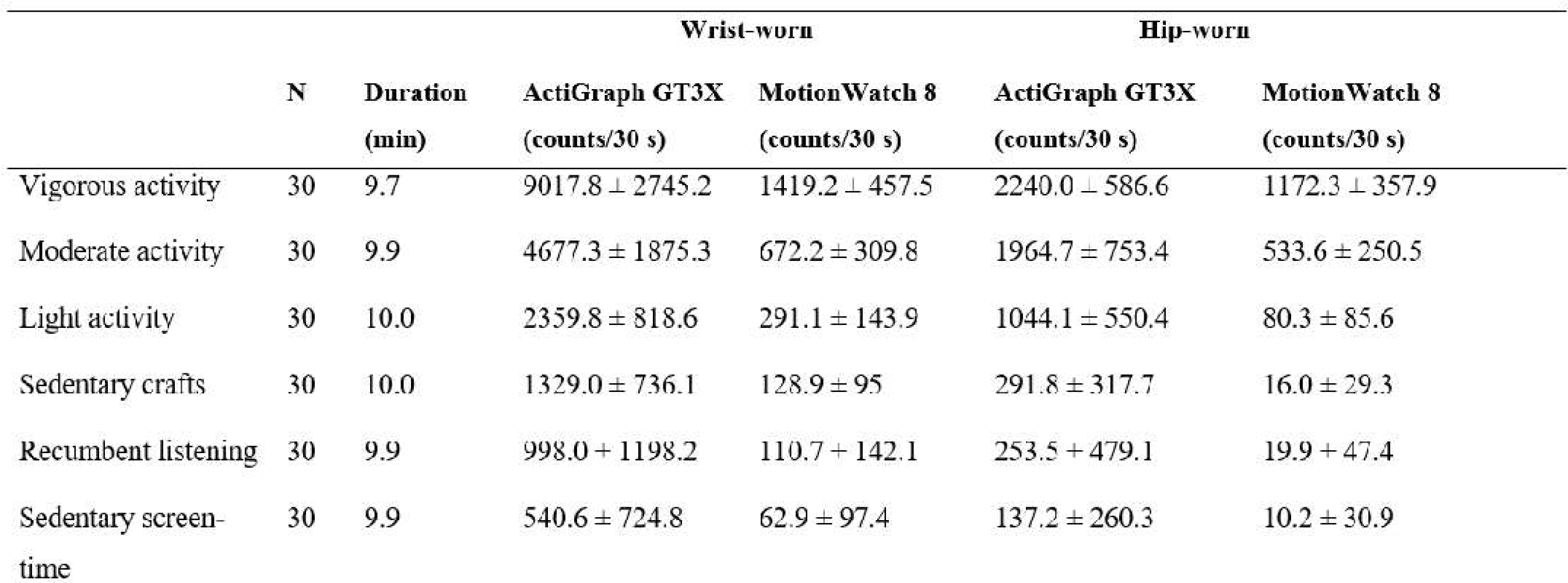
Mean ┴ SD of activity counts (counts/30 seconds) for each activity ranked from highest to lowest intensity measured at wrist-level and hip-level with ActiGraph GT3X and MotionWatch 8. Mean ± SD of the duration spent in activity is consistently similar.

### Comparing MW8 with GT3X revealed significant correlations in activity counts at their respective positions, despite different operating scales

The GT3X’s “vector magnitude” producing proportionally greater values than those of the uniaxis MW8 (**Tab. 2**). When data from the same brand and position were pooled across all activities, significant correlations were detected between counts measured with the MW8 and GT3X for wrist-worn (r = 0.94) and hip-worn devices (r = 0.89) (all p < 0.001, **Fig. 3**). Averaging across all six behaviours, the mean counts/30s were significantly greater for the wrist-position compared to the hip-position within the respective brands (MW8: wrist: 438.6 ± 537 counts versus hip: 300.8 ± 464.3 counts; GT3X: wrist: 3165.3 ± 3357.6 counts versus hip: 959.3 ± 937.8 counts, all p < 0.001). The mean activity counts/30s at wrist- and hip positions simultaneously measured with GT3X and MW8 for each behavioural activity are reported (**Tab. 2**) and visualized as boxplots (**Fig. 4).** The boxplots reveal that the three physically mobile behaviours (light, moderate, vigorous activities) had a narrow within-class variation, enabling to separate each from one another. In contrast, the three physically stationary behaviours (recumbent listening, sedentary crafts, sedentary screen time) showed large interquartile ranges and overlap between classes, regardless of position or brand.

**FIGURE 3.**
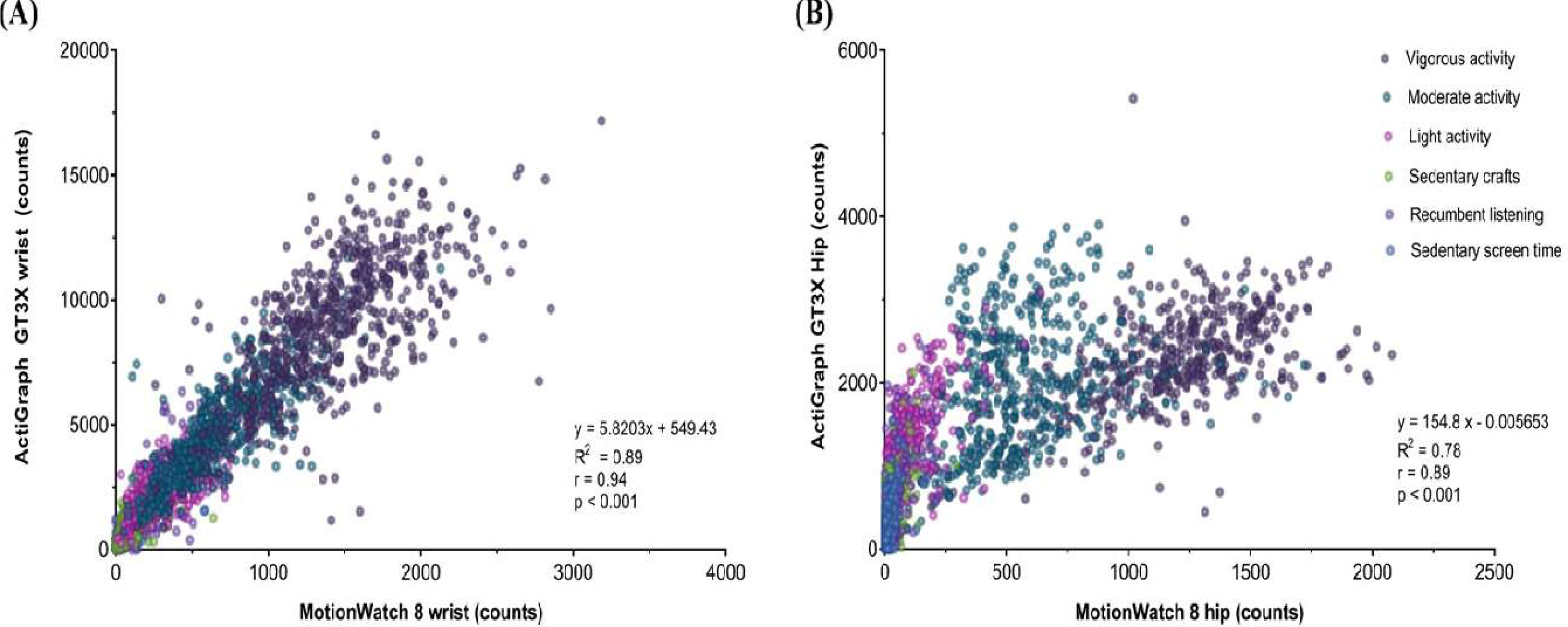
Relationship between activity counts/30s in 3-year-old-children (n=30) during six activities ranging from ‘Motionless alert’ to ‘Vigorous PA’ for Motion Watch 8 (MM8) and ActiGraph (GT3X) at (A) wrist position and (B) hip position. The six activities are colour-coded in the graphs.

**FIGURE 4.**
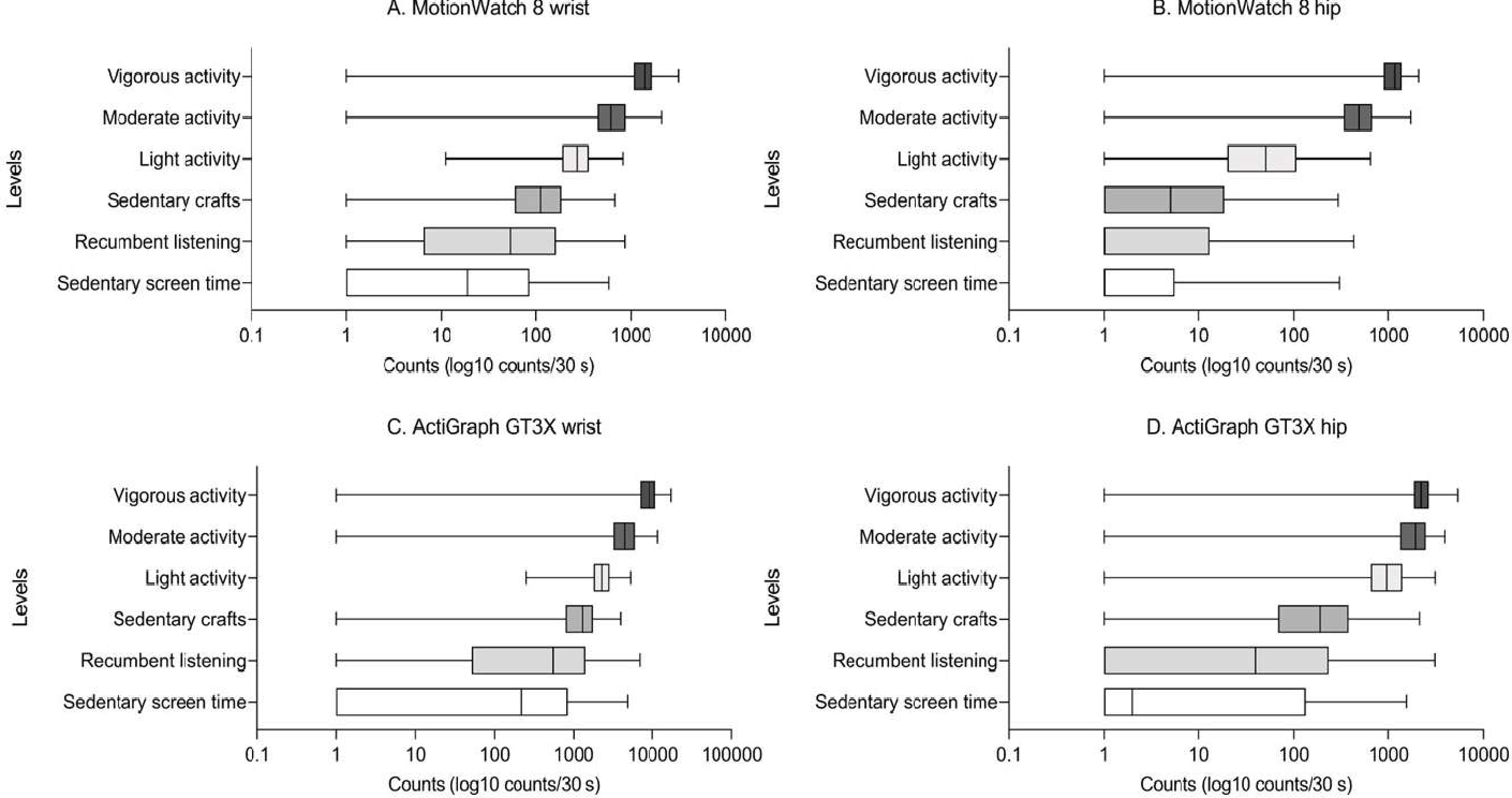
Boxplots for six different intensity levels measured in 3-year-old children with A) wrist-worn MotionWatch 8; B) hip-worn MotionWatch 8; C) wrist-worn ActiGraph GT3X; and D) hip-worn ActiGraph GT3X. Note, a value of ‘I’ was added to every count to increase the visibility of the lower end of the scale, particularly the stationary, calmer activities through a logarithmic scale. Zero movement over 30s epoch is not uncommon, for example during a break. Box contains 50 % of values (Interquaitile range [IQR] with values between Q1 and Q3 (25^th^-75^th^ percentile), mid-line represents median, Whiskers represent min-max ([Q1-1.5*IQR]-Q3+1.5*IQR]).

### Collapsing behavioural classes indicated outstanding accuracy thresholds for ‘mobile PA’ and excellent accuracy thresholds for ‘stationary PA’ intensities

OvR ROC-AUC analysis were carried out four times per device (VPA, MVPA, LMVPA and MOA). Sensitivity ranged from 89 – 95% and specificity from 90 – 94% for MVPA and VPA of wrist-worn devices. Sensitivity for *mobile* PA composite against *stationary* PA was 88% and 91%, and specificity 86% and 83% for wrist-worn MW8 and GT3X, respectively. AUC ranged from 0.95 – 0.98. Sensitivity for MOA (watching cartoon) against the rest was 75% and 77%, and specificity 82% and 85% for wrist-worn MW8 and GT3X, respectively. AUC was 0.86 and 0.87, respectively (**Tab. 3**).

**TABLE 3.** ROC-AUC analysis for wrist-worn MotionWatch 8 and ActiGraph GT3X devices after collapsing categories. See details for collapsed categories in. **Fig. 2**.

|  |  | Wrist-worn MotionWatch 8 (MW8) |  |  |  | Wrist-worn ActiGraph (GT3X) |  |  |  |
| --- | --- | --- | --- | --- | --- | --- | --- | --- | --- |
|  | Intensity class | Cut-off value (counts) | Sensitivity (%) | Specificity (%) | AUC (95% CI) | Cut-off value (counts) | Sensitivity (%) | Specificity (%) | AUC (95% CI) |
| Mobile PA | VPA<sup>1</sup> | >787 | 93 | 94 | 0.98<br>(0.97 to 0.98)* | >4607 | 95 | 90 | 0.97<br>(0.97 to 0.98)* |
|  | MVPA<sup>2</sup> | >408 | 90 | 93 | 0.97<br>(0.97 to 0.98)* | >3038 | 89 | 93 | 0.97<br>(0.96 to 0.98)* |
|  | LMVPA<sup>3#</sup> | >215 | 88 | 86 | 0.95<br>(0.94 to 0.96)* | >1782 | 91 | 83 | 0.95<br>(0.94 to 0.95)* |
| Stationary PA | SED<sup>4</sup> | 118-215 | NA | NA | NA | 1148-1782 | NA | NA | NA |
|  | MOA<sup>5</sup> | <118 | 75 | 82 | 0.86<br>(0.84 to 0.87)* | <1148 | 77 | 85 | 0.87<br>(0.86 to 0.89)* |

\* p < 0.001
**#LMVPA: separates mobile physical activities (Mobile PA) from stationary activities (Stationary PA)**
<sup>1</sup>VPA: Vigorous physical activity versus all other activity levels
<sup>2</sup>MVPA: Moderate-vigorous physical activity versus all other activity levels (light, sedentary [handcraft + recumbent], motionless)
<sup>3</sup>LMVPA: light moderate vigorous physical activity versus the rest (sedentary [handcraft + recumbent], motionless)
<sup>4</sup>SED: Sedentary range: above MOA threshold and below LMVPA threshold
<sup>5</sup>MOA: Motionless-alert [Screen time] versus all other activity levels

Sensitivity ranged from 91 – 96%, and specificity from 81 – 98% for MVPA and VPA of hip-worn devices. Sensitivity for *mobile* PA composite against *stationary* PA was 85% and 91%, and specificity 91% and 89% for hip-worn MW8 and GT3X, respectively. AUC ranged from 0.90 – 0.98. Sensitivity for MOA (watching cartoon) against the rest was 64 and 75%, and specificity 87 and 78% for hip-worn MW8 and GT3X, respectively. AUC was 0.81 and 0.86, respectively (**Tab. 4**).

**TABLE 4.**
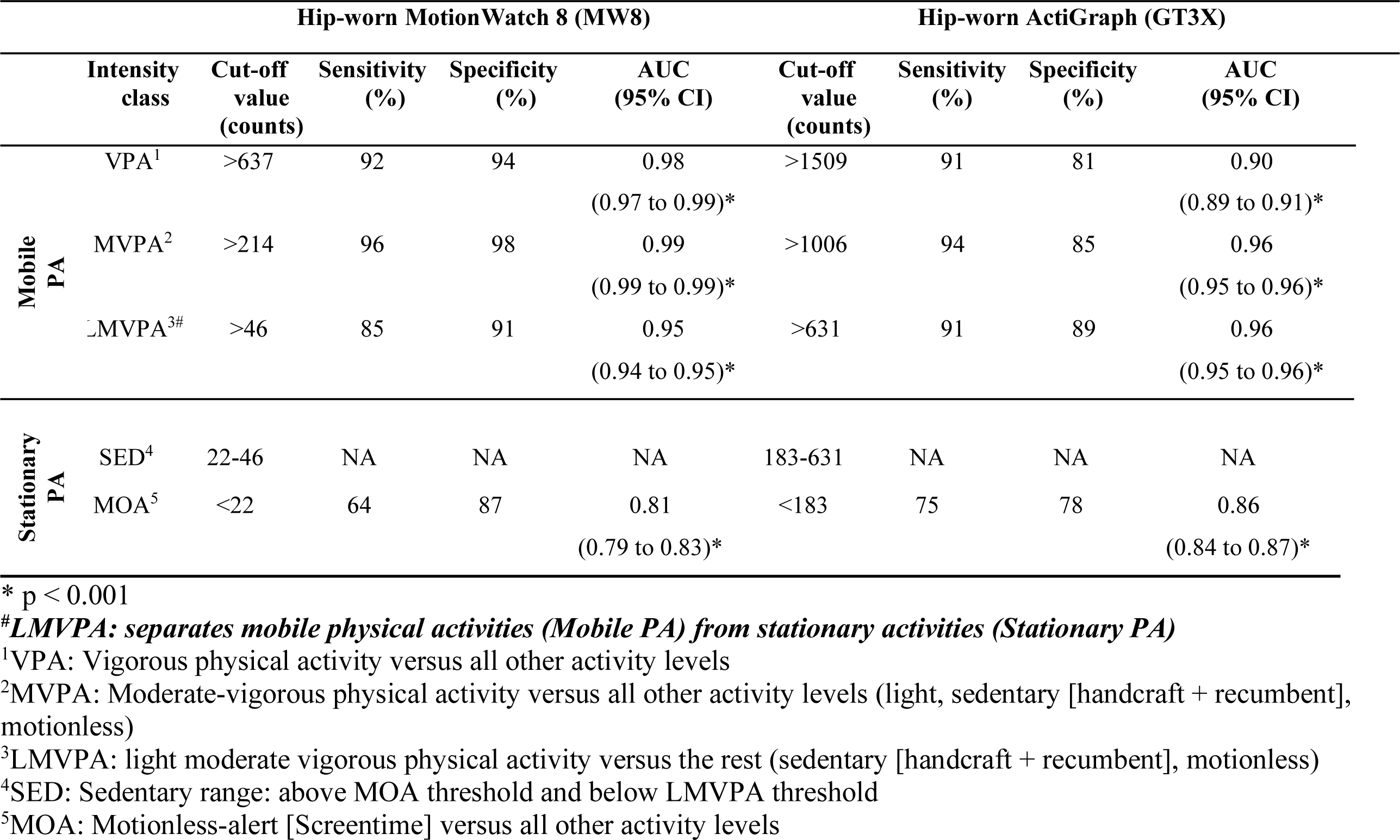
ROC-AUC analysis for hip-worn MotionWatch 8 and ActiGraph GT3X devices after collapsing categories. See details for collapsed categories in. **Fig. 2**.

## DISCUSSION

This study aimed at providing calibration data for 3-year-old children performing various play activities and passive behaviours by means of count cut-off thresholds from wrist-worn MW8 and GT3X activity counts/30s. Additional purposes included comparison of classification accuracy between concomitantly worn wrist- and hip-mounted devices using sensitivity, specificity and ROC-AUC. No previous studies have developed activity intensity thresholds for wrist- and hip-worn GT3X models at 30s-epochs in vector mode (VM). But thresholds have been published for GT3X models at 5s-epochs using a sampling rate of 30Hz and wrist/hip positions. The observation technique (video during free play of groups of children) and the ‘Children’s Activity Rating Scale’ (57, 58) differed, and as a result observations included some similar (watching cartoons, drawing) as well as different activities (dancing, outdoor) (12). Compared to their cut-offs, the intensity thresholds established for GT3X in VM in the present study were higher. This was expected from an epoch five times as long, confirming thresholds to be epoch-specific in addition to being age-specific (12).

We found substantial overlap in activity counts among the three physically *stationary’* activities as evident from the sizable variation in the interquartile range of the box plots. Already two decades ago, correlations between activity counts and oxygen consumption were reported for rest/structured activities and free play (r = 0.82 and r = 0.66, respectively) in 3-5-year old children (13), whereupon stationary behaviours, also termed ‘physical *inactivity’* became recognised as a distinct construct (59). The Children’s Physical Activity Research Group at the University of South Carolina has made significant contributions to better understand physical activity behaviours in children (21, 60). Among those, most notably the development of observational instruments to assess PA context-specific, and the implementation of accelerometry cut-off points in settings of different age groups. This qualitative-quantitative combination can be applied in longitudinal actigraphic recordings that include naps and nocturnal sleep in their recordings. The current study adopted the term ‘motionless alert’ instead of using ‘rest/resting’ because the term ‘rest’ is inconsistently used (61). In chronobiology, ‘rest’ covers ‘sleep’ in rest-activity rhythms and is used to describe ‘time-in-bed’ that includes ‘latency to sleep’ (transitional stage) and ‘sleep period’ (motionless body with an active brain), specifically in connection with longitudinal actigraphy / accelerometry data. The Sedentary Behaviour Research Network has published consensus definitions to standardise terminology for sedentary behaviours using ‘posture’ information and ‘energy expenditure’ (62). They also adopted the term ‘stationary’ and suggested to apply ‘behaviour’ when the context is known, and ‘time’ in the absence of context, which is a sensible linguistic differentiation.

Our cut-off points corroborate the distinction between physically *mobile* behaviours and physically *stationary* behaviours, which is in line with results of earlier studies, despite using different measures and criteria of classifying physical activity intensities (10, 12, 32, 47).

The movement intensities differed not just by behaviour but also by type of device (different sensors, algorithms) and body position (wrist, hip). Nevertheless, the count cut-offs from both devices (MW8, GT3X) at both positions (wrist, hip) showed outstanding accuracy (range: 0.95-0.98) for all *mobile* PA classes. The *mobile* PA classes of wrist-worn MW8 cut-offs showed specificities of 94%, 93% (VPA, VMPA, respectively) and 90% and 93% for the wrist-worn GT3X. Separating all *mobile* PA classes from all *stationary* PA classes combined yielded specificities of 86% for wrist-worn MW8 cut-offs and 83% for wrist-worn GT3X cut-offs. An important objective of observing the behaviour while measuring movement intensities was to test the discrimination power for motionless alert behaviour, such as watching a cartoon, from all other classes. Although acceptable, ‘motionless alert’ from the wrist-worn MW8 and GT3X cut-offs showed only fair sensitivity (75% and 77%), while sensitivity was even lower for hip-worn cut-off points. The classification accuracy ranged between 0.81-0.87. The better outcome of the *mobile* PA classes over *stationary* PA classes can be explained by greater density of counts with much narrower ranges due to more steady, continuous activity during ‘sprinting, ‘floor ball’ and ‘playing on the floor’. In comparison, the *stationary* PA classes, and motionless alert in particular, included children engaged in mental tasks (listening, concentrating, observing) with the occasional, discontinued burst of movements while sitting still, for example when pointing or changing of position. To distinguish motionless alert from sleep, diary entries and time pattern analyses would be helpful to consult.

The purpose of conducting this calibration study was to develop count cut-off points to be implemented in a child birth cohort. In direct comparison, the two device models systematically scaled count quantities differently irrespective of unequal movements of limbs and trunk (wrist/hip). Since all manufacturers calibrate their models at factory for optimal recording performance suited for certain body position, there is an expected imbalance in counts related to body positions. For example, the MW8 is intended to monitor limb or body movements during daily living with light exposure (in lux) when attached to the wrist and also to calculate sleep parameters from rest-activity time patterns. It offers various recording options such as uni-axial mode that has been calibrated for wrist position for high compliance in long-term recordings, making use of omni-rotary arm movements and light capture (**Tab. S1**) (63). Its tri-axial mode produces VM/epochs that is best suited for the trunk, i.e., the hip, but not for measuring sleep nor light (64). We mounted a second MW8 in uni-axial mode at the hip position to see how the same mode scales wrist against hip position, accepting its compromised status of reduced resolution. Lower activity counts were found for all behavioural activities at hip position compared to the wrist. Between brands, the GT3X model, which was set to tri-axial mode as recommended, generated approximately 6 to 10 times higher values in VM mode than the uni-axial MW8 mode, regardless of position. The challenge with recording and processing raw 3-dimensonal *longitudinal* data is their volume - in the TeraByte range - and the need to develop approaches on how to visualise 3-dimensional patterns, most summarise them into a single vector/epoch (65). Users need to be aware of these facts and decide accordingly on sensor modes, pre-filter/raw settings, algorithms when implementing activity monitoring into longitudinal recordings.

The MW8 has previously been validated to measure physical activity in older adults (**Tab. S1**) (45) and in 9-13 year old children (47). Lin et al’s (45) protocol differed from our study in that they captured data from the dominant wrist position of older children. We preferred the non-dominant wrist position because it has been shown to produce less misclassification for sedentary behaviours (41). Despite the differences, our results are in line for *mobile* PA activities with Lin et al. (47), who reported a good ability of the device to differentiate light from moderate-to-vigorous activity (MVPA >371.5 counts/30 s), and moderate from vigorous activity (VPA >859.5 counts/30 s). Our boundaries for those categories are a little lower (MVPA >216 counts/30s; VPA >788 counts/30s), likely as a result of age-related differences in motor development and muscle strength. In Johansson et al. (12), authors argued for age-specific thresholds since they found that 4-year-old preschool children, who wore a GT3X triaxial on their non-dominant wrist, produced generally higher intensity thresholds per 5s-epoch in comparison to toddlers of 2 years (19). Differentiation of sedentary from light activity was much greater in the study by Lin et al (45), given the much lower sedentary threshold score (SED < 32 counts/30s) compared to the present study (SED < 215 counts/30s, MOA < 118 counts/30s). This is likely related to Lin et al (45) prioritizing higher specificity at the cost of lower sensitivity in order to minimise false positives due to arm movements. Johansson et al (12) used the opposite adjustment - towards higher sensitivity at the expense of a lower specificity for the SED threshold in order to avoid underestimating the time spent in SED. In the present study we used a step-approach with the upper boundary for SED being the lower boundary of *mobile* PA classes against all combined *stationary* PA classes, which includes arm movements, i.e. when sitting and doing crafts. All threshholds were reported as the greatest sum of the sensitivity and specificity, without prioritising specificity nor sensitivity.

### Strength and limitations

One of the strengths is the study design that included two research-grade devices (MW8, GT3X) worn in parallel at two body positions (wrist and hip) in a young age group (3 years). In addition, the use of a 30 second epoch matches the calibration of MW8 devices for sleep assessments and meets the requirements for longitudinal, long-term actigraphic data collection for circadian rhythm/sleep assessments. The present calibration allows PA analysis from measurements of physical activity in combination with circadian/sleep assessments. The purpose of it is a multi-modal outcome with relevant information for trajectories of 24-h sleep and PA sufficiency to to be comparable and interpretable with other studies and contribute to charts of 24h sleep and PA percentile curves. Half a minute may seem very long for sport activities but human behaviours under real situations last for several minutes, in particular conversations, meal and play times, sedentary activities like handcrafts, drawing or screen time. On the other hand, vigorous activities over the sequence of 10 minutes were not uninterrupted. Instead, there were periods of short light level movements.

Another strengths is the application of a commonly used classification method on directly observed behaviours to quantify intensity levels from which count cut-off points were determined. The intensity thresholds were chosen based on the best compromise between sensitivity and specificity, since prioritising sensitivity for ‘physically stationary’ may inflate sedentary time, when it is actually time spent in light mobile PA. Prioritsing specificity for stationary PA may overestimate ‘physically mobile’ and underestimate time spent sedentary. Our approach is a compromise as are those of others (5). Our calibration may be applicable to existing population datasets since it allows to derive scaling coefficients. When other studies used 5 second epochs these can be re-scaled to match the 30s epoch for comparison, but this conversion needs verification as deliniated by Orme et al (66). The dataset generated during the current study is available at ‘Behaviour-based movement calibration’ (https://www.katlab.org/people/).

The absence of indirect calorimetry (67) could be seen as a limitation, but oxygen consumption has been measured indirectly in 4-year old preschool children and activity intensity levels at free play correlated well with oxygen consumption, yet it has been discussed to have limitations due to the delayed rate of change with a change in activity levels (13). We are aware that our thresholds have not undergone cross-validation and all of the observed activities were performed indoors, although resembling everyday habits of a three-year-old child. Outdoor activities were deliberately left out because of the high seasonal variability in outdoor activities in Northern Sweden. Further work is required to examine season-based behavioural outdoor activity patterns.

## CONCLUSION

MotionWatch 8 and the ActiGraph GT3X underwent calibration processing simultaneously on the non-dominant wrist and hip based on direct observation of six naturally-occuring behavioural activities in healthy, 3-year-old children. ROC-AUC curves revealed cut-off points with outstanding, excellent and good discrimination power. The accuracy indicates that the calibrated cut-off points can be adequately used to determine time spent in different activity levels, under the provision that the data match the follwing calibration factors (i) age, (ii) the device-specific sensor mode, (iii) epoch (scaling coefficient, if different), (iv) body position, and (v) behaviours equivalent to occurring in natural settings. The dataset will be made freely available, which is hoped to be useful for the community interested in making decisions on integrating physical activity intensities into projects on sleep, circadian rhythms and light exposure from a 24-hour movement-related perspective.

## Author contributions

Conceptualization: KW MD JS; Data curation: KW MD; Methodology: KW MD JS DJ; Formal analysis: RW DJ MD KW; Resources: MD CW KW JS; Writing - original draft: RW DJ KW; Writing - review & editing: KW MD DJ CW JS; Supervision: KW MD; Funding acquisition: MD KW JS CW.

## Supporting information

WULFF Supplementary Material

## Acknowledgements and Disclosures

The authors thank all children and their parents for their participation. Also, we thank Anna Tellström and Rebecca Rönnholm for their great assistance in the data collection; Tobias Stenlund for all organisational help with the e-health laboratory; Patrik Wennberg for lending us the two Actigraph GT3X devices and the accompanying temporary license; and Richard Lundberg for his administrative support with the NorthPop database.

## Conflict of interest

The authors have no conflict of interest to declare. The results of the study are presented clearly, honestly, and without fabrication, falsification, or inappropriate data manipulation.

### SUPPLEMENTARY DIGITAL CONTENT

ACTIGRAPHY OPERATIONAL MANUAL

https://www.katlab.org/wp-content/uploads/2023/10/ACTIGRAPHY-OPERATIONAL-MANUAL-Nordic-Daylight-Research-Programme-2023.pdf

RAW DATA EXCEL SPREADSHEET

Access via ‘Behaviour-based movement calibration’ excel spreadsheet at https://www.katlab.org/people/ under Networking and Funding.

## Data availability statement

The data supporting the findings in the study are freely available through an URL link in this article.

## Ethical approval statements

The study was conducted in agreement with the declaration of Helsinki and approved by the regional ethical review board (Dnr 2020/01254).

