## Supplementary material for "Behaviour-based movement cut-off points in 3-year old children comparing wrist- with hip-worn actigraphs MW8 and GT3X": WULFF Supplementary Material

Daniel Jansson[1, 2], Rikard Westlander[3], Jonas Sandlund[4], Christina E. West[3], Magnus Domellöf[3#], Katharina Wulff[5, 6,#,*]

Daniel Jansson[1, 2] (ORCID ID 0000-0002-6488-0663)
Rikard Westlander[3] (ORCID ID 0000-0002-7874-4320)
Jonas Sandlund[4] (ORCID ID 0000-0001-5403-881)
Christina E. West[3] (ORCID ID 0000-0001-9599-2580)
Magnus Domellöf[3] (ORCID ID 0000-0002-0726-7029)
Katharina Wulff[5, 6] (ORCID ID https://orcid.org/0000-0003-2480-3329)

[1] Department of Community Medicine & Rehabilitation, Section of Sports Medicine, Umeå University, Umeå, Sweden
[2]Umeå School of Sport Sciences, Umeå University, Umeå, Sweden
[3]Department of Clinical Sciences, Pediatrics, Umeå University, Sweden
[4]Department of Community Medicine and Rehabilitation, Section of Physiotherapy, Umeå University, Umeå, Sweden
[5]Departments of Radiation Sciences and Molecular Biology Umeå University, Umeå, Sweden
[6]Wallenberg Centre for Molecular Medicine (WCMM), Umeå University, Umeå, Sweden

**# Joint senior authors.**

Department of Molecular Biology, 6L, Sjukhusområdet, Umeå universitet, 901 87 Umeå, Sweden.

06 January 2024

### Overview


- Method utilisation: Accelerometry and Actigraphy …..…..………2
- ROC curve results of adjacent One-vs-One ROC …………………3
- Figures of One-versus-Rest ROC results in main text……..………5
- Manual and data availability………………………………………9
- References…………………………………………………….9


#### Methods utilisation

Sensors can be used for many purposes. In science the important decision that must be made is: "Which methods are the most appropriate ones to answer the research question?"

Perhaps unsurprisingly, different disciplines have utilised activity monitoring differently to address their research questions. **Table S1** highlights some examples: Sports medicine has been interested in capturing PA intensities only during daytime, often to be used as a surrogate for energy expenditure. Chronobiology on the other hand utilised the monitoring technique to capture the patterns of rest and activity timing during daytime and during sleep at night-time. Currently, the trend is to make use of both approaches in a complementary way to analyse behaviours over 24-h cycles and across weeks.

**Table S1**. Highlighting some of the different approaches taken from sports medicine and chronobiology/sleep, with selected published examples.

| **Sports medicine** | **Examples** | **Sleep / Chronobiology** | **Examples** |
|---|---|---|---|
| Focus on physical activity alone | (1, 2, 10, 28) | Focus on rest-activity cycles and sleep | (3, 4, 18, 34) |
| Accelerometry | (12, 17, 24) | Actigraphy / Actimetry | (15, 36, 37) |
| Only short, day-time physical activity | (39, 40, 42) | Long-term, 24-h movements over time | (19, 26, 34) |
| Placement mainly on hip/waist | (14, 16, 30, 38) | Mainly on non-dominant wrist | (16, 23, 26, 41) |
| Epoch max. 15 seconds | (5, 6, 7, 8, 9, 10, 11) | Epoch 30 – 60 seconds | (19, 26, 36) |
| Validation against energy expenditure | (10, 25, 31) | Algorithms validated against polysomnography | (18, 20, 21, 22) |
| Defining activity intensity levels | (13, 14) | Defining temporal patterns and sleep parameters | (15, 35) |
| **Comparison between positions and brands** | | | |
| (27, 29, 32, 33) | | | |

#### ROC curve results of One-vs-One ROC and Wear-time

The first round of ROC analysis was performed based on the One-vs-One (OvO) scheme (**Table S2)**. Cut-off points were derived from pairwise combination of counts of adjacent behavioural activities. The behavioural activities were sorted from highest to lowest expected intensities to evaluate the accuracy of separating thresholds stepwise from high physical behaviour (sprinting) to motionless-alert (watching cartoons), see schematic (**Fig. 2a** in the main text).

**Table S2.** Receiver operating characteristics curve (ROC) analysis in One-vs-One scheme, comparing pairwise combination of adjacent behavioural classes as described in **Fig. 2a.** Cut-off values, sensitivity, specificity and accuracy (AUC) are reported from 'vigorous' to 'motionless alert'. 'Sedentary crafts' and 'Recumbent listening' were merged into 'Sedentary (active)'. 'Sedentary screen time is categorised as 'Motionless alert'.

| | **Wrist- worn MotionWatch 8** | | | | **Wrist-worn ActiGraph (GT3X)** | | | |
|---|---|---|---|---|---|---|---|---|
| | Cut point value (counts) | Sensitivity (%) | Specificity (%) | AUC (95% CI) | Cut point value (counts) | Sensitivity (%) | Specificity (%) | AUC (95% CI) |
| Vigorous activity[1] | >1040 | 80.4 | 87.6 | 0.91 (0.89 to 0.93)* | ≥6677 | 81.7 | 86.0 | 0.90 (0.88 to 0.92) * |
| Moderate activity[2] | >445 | 76.6 | 87.4 | 0.88 (0.86 to 0.90) * | >3402 | 71.9 | 90.6 | 0.87 (0.85 to 0.89) * |
| Light activity[3] | >154 | 84.9 | 70.7 | 0.85 (0.82 to 0.87) * | >1760 | 79.0 | 77.2 | 0.84 (0.82 to 0.86) * |
| Sedentary (active) | 24.5-154 | NA | NA | NA | 933-1760 | NA | NA | NA |
| Motionless alert[4] | <24.5 | 76.8 | 53.4 | 0.69 (0.66 to 0.71) * | <933 | 54.0 | 80 | 0.71 (0.69 to 0.71) |

| | **Hip-worn MotionWatch 8** | | | | **Hip-worn ActiGraph (GT3X)** | | | |
|---|---|---|---|---|---|---|---|---|
| | Cut point value (counts) | Sensitivity (%) | Specificity (%) | AUC (95% CI) | Cut point value (counts) | Sensitivity (%) | Specificity (%) | AUC (95% CI) |
| Vigorous activity[1] | >875 | 82.9 | 92.2 | 0.92 (0.90 to 0.93) * | >1866 | 77.3 | 46.2 | 0.62 (0.59 to 0.65) * |
| Moderate activity[2] | >230 | 92.2 | 94.2 | 0.98 (0.97 to 0.99) * | >1655 | 66.1 | 87.8 | 0.84 (0.81 to 0.86) * |
| Light activity[3] | >13 | 83.3 | 71.0 | 0.84 (0.81 to 0.86) * | >442 | 86.9 | 81.3 | 0.90 (0.88 to 0.91) * |
| Sedentary (active) | 2-13 | NA | NA | NA | 16-442 | NA | NA | NA |
| Motionless alert[4] | <2 | 47.6 | 69.6 | 0.59 (0.56 to 0.620) | <16 | 76.7 | 58.4 | 0.68 (0.65to 0.70)* |

*p< 0.001

[1]Vigorous activity versus moderate activity

[2]Moderate activity versus light activity

[3]Light activity versus merged sedentary crafts and recumbent listening

[4]Merged sedentary crafts and recumbent listening versus sedentary screen time

The OvO ROC-AUC analysis returned cut-offs of good to excellent probability of correctly identifying adjacent '*physically mobile'* classes: 'vigorous', 'moderate' and 'light' activity (**Tab. S2**). However, the OvO ROC-AUC analysis returned cut-offs of fair probability of correctly identifying adjacent '*physically stationary'* behaviours for the GT3X devices and cut-offs of poor probability for the MW8 devices, regardless of position (**Tab. S2**). The large interquartile ranges and great overlap between classes among '*physically stationary'* activities, especially 'sedentary screen time' and 'recumbent listening', likely precluded reliably excluding counts not belonging to their respective class (see **Fig. 4,** in the main text).

#### Wear-time

The total wear time for each behavioural class, summed up across all participants (N=30) per class showed a very similar wear time for every position (**Fig. S1**). The duration to perform each behaviour was set to 8-10 min to allow 16 to 20 epochs per person to get a good capture of the within-person's spread of intensities performing these behaviours. Start and stop time was directly observed with a stopwatch. This resulted in total to about 300 min per behaviour and about 600 activity counts per behaviour for the analysis.

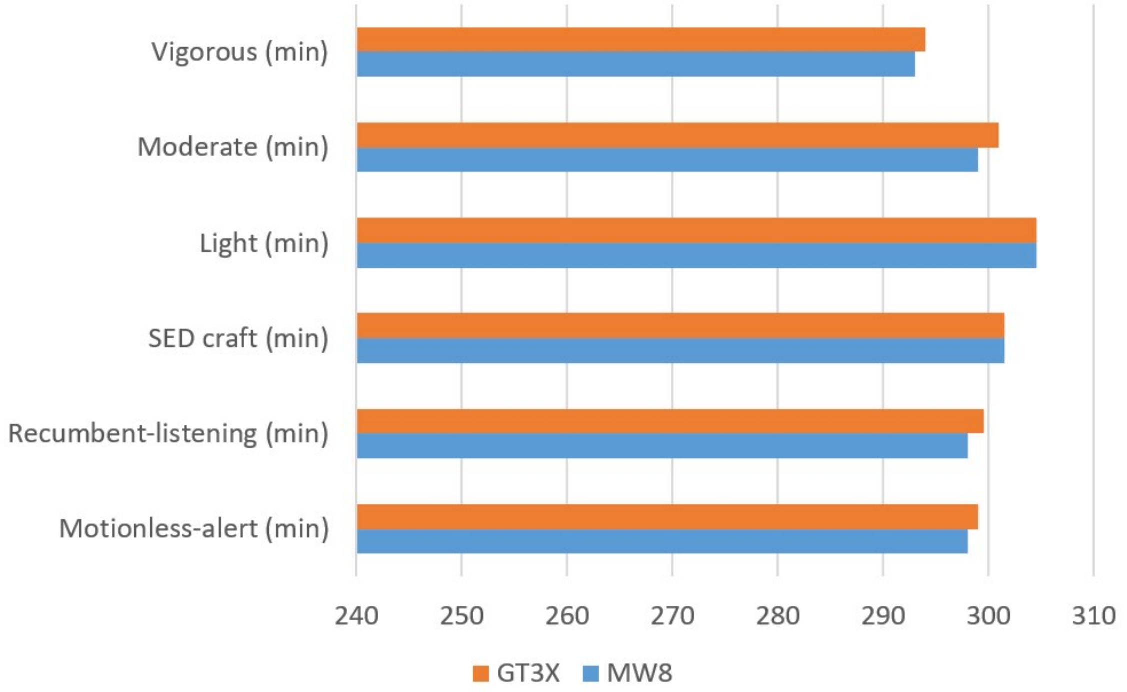


**Fig. S1**. Duration in minutes of cumulated wear-time of devices (MW8, GT3X) simultaneously worn at wrist and hip positions while performing each of the six behaviours. The epoch for storing the activity counts was 0.5 min.

**Figures of One-versus-Rest ROC curves supplementing the results in the main text**

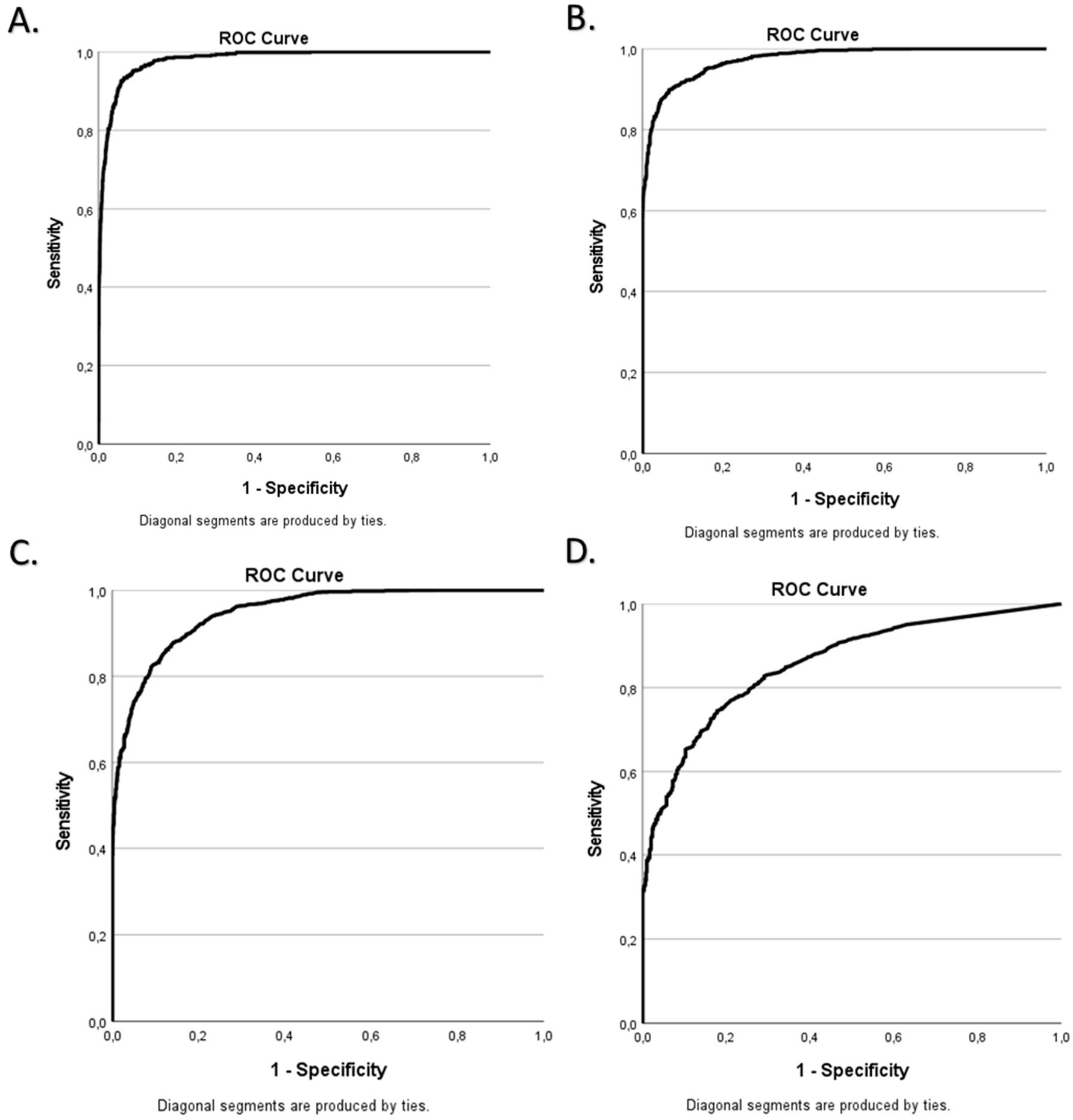


**Fig. S2 ROC curves for the wrist-worn Motionwatch 8.** OvR ROC curves illustrating the results in table 3 and table 4 in the main text for: **A**) vigorous physical activity (VPA), **B)** moderate-vigorous physical activity (MVPA), **C)** light moderate vigorous physical activity (LMVPA) and **D**) motionless-alert (MOA).

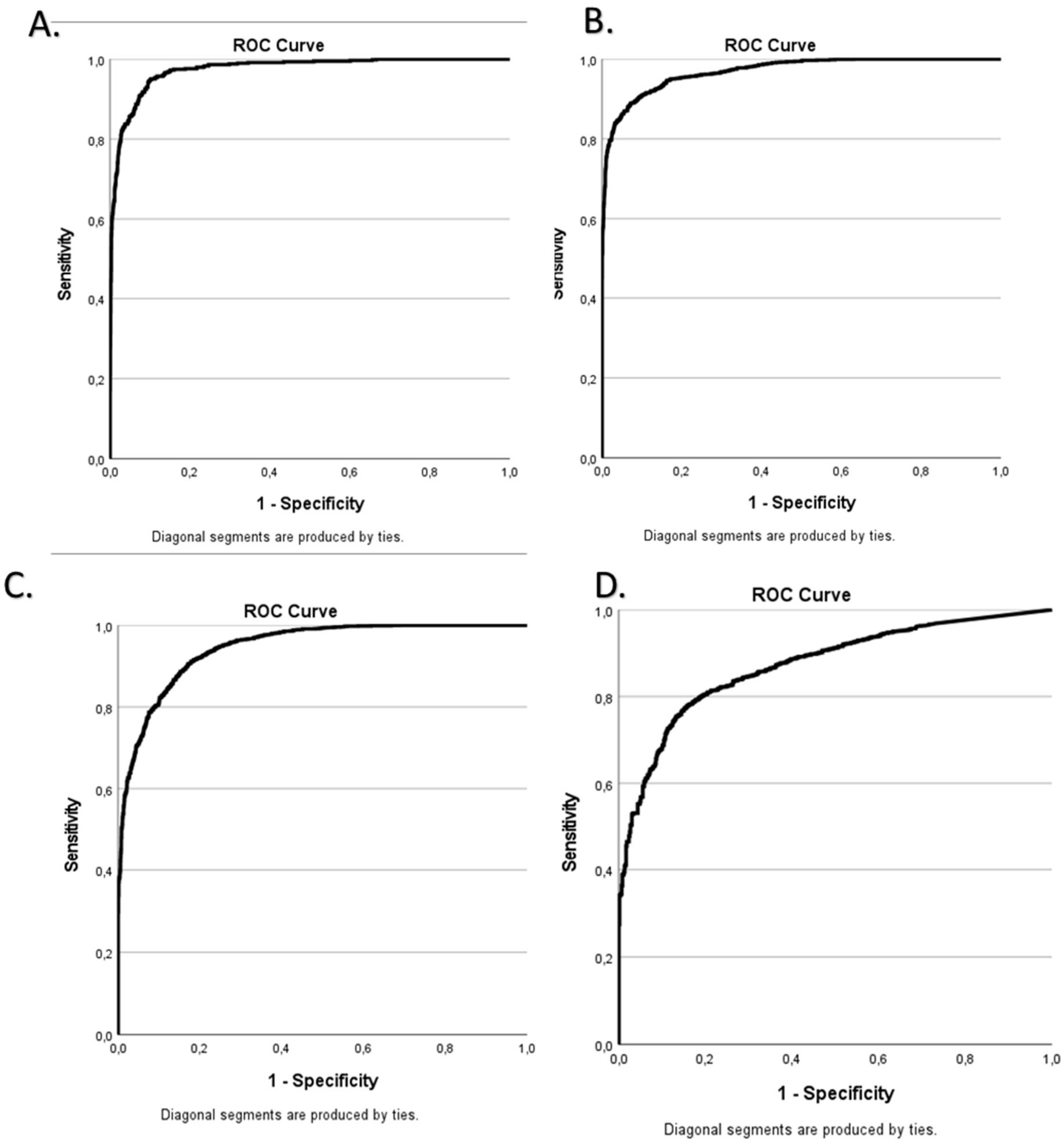


**Fig. S3 ROC curves for wrist-worn GT3X.** OvR ROC curves illustrating the results in table 3 and table 4 in the main text for: **A**) vigorous physical activity (VPA), **B)** moderate-vigorous physical activity (MVPA), **C)** light moderate vigorous physical activity (LMVPA) and **D**) motionless-alert (MOA).

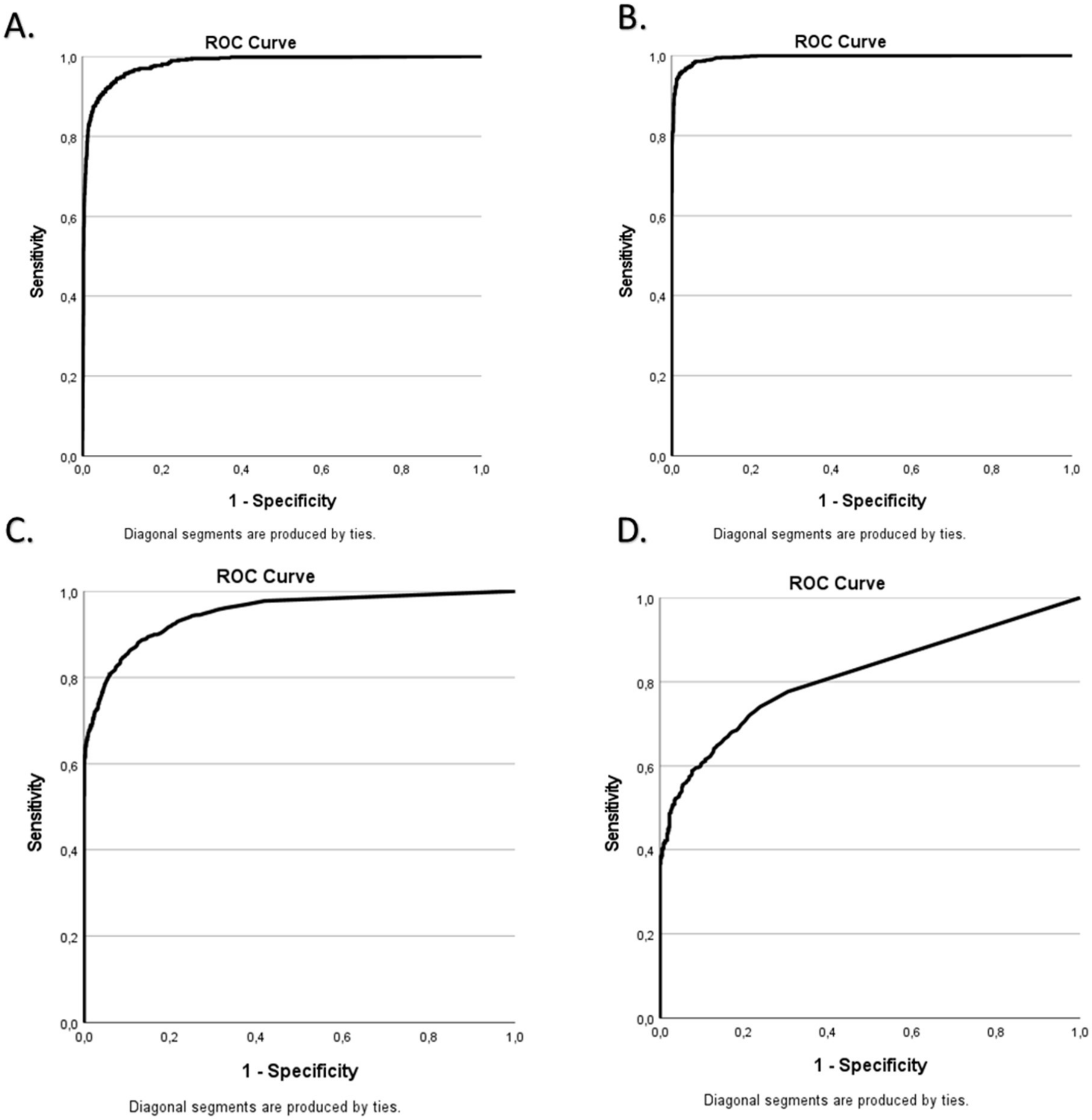


**Fig. S4 ROC curves for hip-worn Motionwatch 8.** OvR ROC curves illustrating the results in table 3 and table 4 in the main text for: **A**) vigorous physical activity (VPA), **B)** moderate-vigorous physical activity (MVPA), **C)** light moderate vigorous physical activity (LMVPA) and **D**) motionless-alert (MOA).

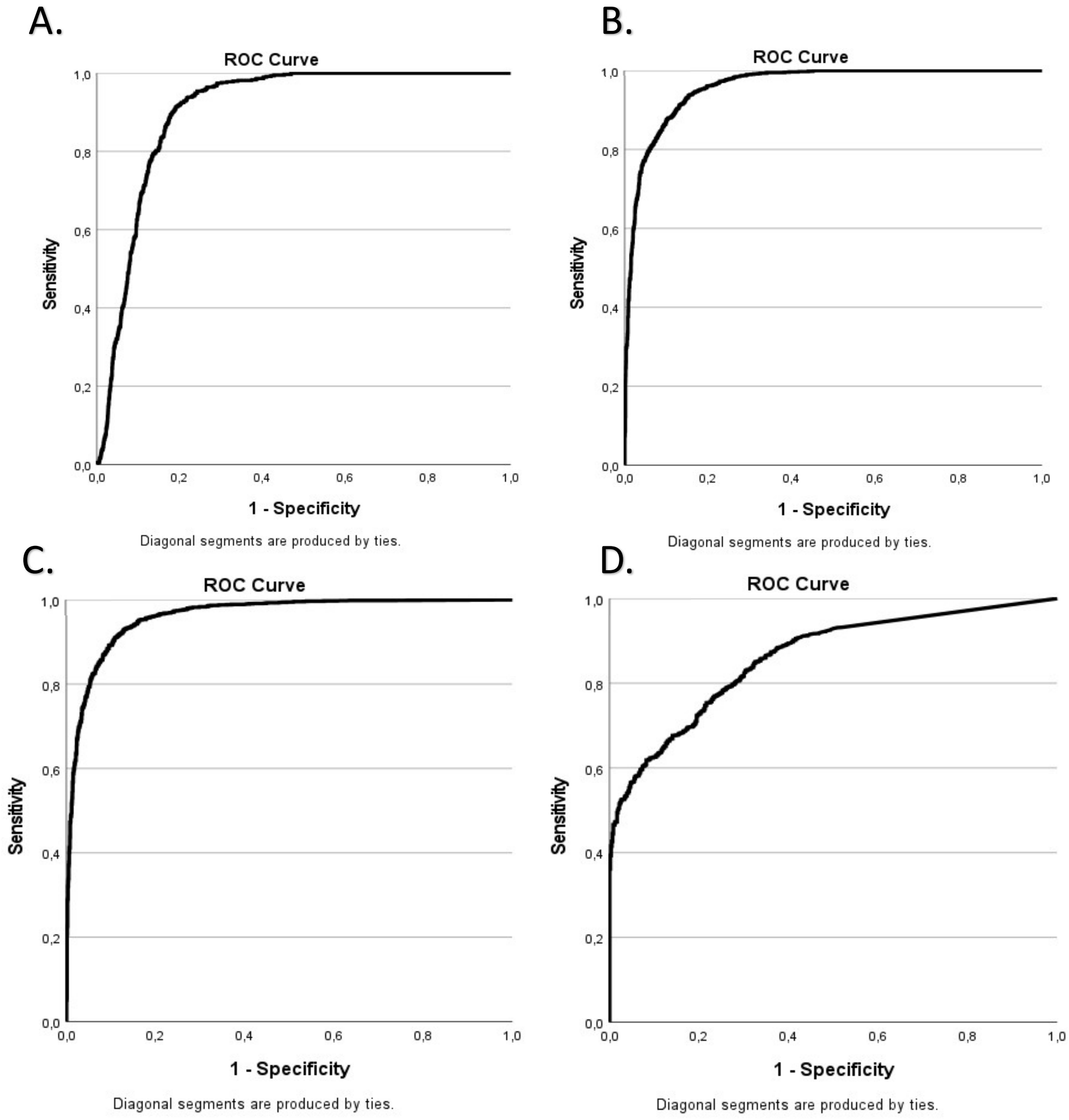


**Fig. S5 ROC curves for hip-worn GT3X.** OvR ROC curves illustrating the results in table 3 and table 4 in the main text for: **A**) vigorous physical activity (VPA), **B)** moderate-vigorous physical activity (MVPA), **C)** light moderate vigorous physical activity (LMVPA) and **D**) motionless-alert (MOA).

#### Manual and data availability

| | |
|---|---|
| Data availability | The data that supports the findings of this study are available from https://www.katlab.org/people/<br>Under 'Networking and Funding':<br>LINK: Behaviour-based movement calibration |

| | |
|---|---|
| Manual availability | The manual that is used for data collection and data analysis is available from https://www.katlab.org/people/<br>Under 'Networking and Funding': Manuals |

#### References (Supplementary to Table S1)

35. Skeldon AC, Dijk DJ, Meyer N, Wulff K. Extracting Circadian and Sleep Parameters from Longitudinal Data in Schizophrenia for the Design of Pragmatic Light Interventions. Schizophr

Bull. 2022 Mar 1;48(2):447-456. doi: 10.1093/schbul/sbab124. PMID: 34757401; PMCID: PMC8886588.
